# oFVSD: A Python package of optimized forward variable selection decoder for high-dimensional neuroimaging data

**DOI:** 10.1101/2022.12.25.521906

**Authors:** Tung Dang, Alan S. R. Fermin, Maro G. Machizawa

## Abstract

Neuroimaging data is complex and high-dimensional that poses challenges for machine learning (ML) applications. Of varieties of reasons contributing on accuracy decoding, variable feature selection is one of crucial steps for determining target feature in data analysis, especially in the context of neuroimaging studies where the number of features is often much larger than the number of observations. Therefore, optimization of feature selection from such high-dimensional neuroimaging data has been challenging using conventional ML algorithms. Here, we introduce an efficient ML package incorporating a forward variable selection (FVS) algorithm that optimizes the identification of features for both classification and regression models. In our framework, the best ML model and feature pairs that explain the inputs can be automatically determined. Moreover, the toolbox can be executed in a parallel environment for efficient computation. The parallelized FVS algorithm iteratively selects the best feature pair compared against the previous steps to maximize the predictive performance. The FVS algorithm evaluates the goodness-of-fit across different models using the k-fold cross validation and identifies the best subset of features based on a pre-defined criterion for each model. Furthermore, the hyperparameters of each ML model are optimized at each forward iteration. A final outcome highlights an optimized number of selected features (brain regions of interest) with decoding accuracies. Using our pipeline, we examined the effectiveness of our toolbox on an existing neuroimaging (structural MRI) dataset. Compared ML models with and without FVS approach, we demonstrate that the FVS significantly improved the accuracy of the ML algorithm over the counterpart model without FVS. Furthermore, we confirmed the use of parallel computation considerably reduced the computational burden for the high-dimensional MRI data. This oFVSD toolbox efficiently and effectively improves the performance of both classification and regression models on neuroimaging data and should be applicable to many other neuroimaging data and more. This Python package is open-source and freely available, making it a useful toolbox for neuroimaging communities seeking improvement of decoding accuracy for their datasets.

## 1. Introduction

Neuroimaging data such as structural and functional magnetic resonance imaging (MRI) data provide information about neuronal structure and activity with relatively high spatial resolution, and play an important role in providing researchers with unprecedented access to the inner workings of the brain of a healthy individual or an individual with a neurological disease or psychiatric disorder. Identifying brain regions that can be used to differentiate healthy and nonhealthy participants or predict changes in patients’ pathological states and remission-levels are some of the most important purposes of neuroscientific studies.

There is an increasing number of studies turning to ML approaches to extract exciting new information from neuroimaging data. For example, the partial least squares (PLS) approach was proposed to extract distributed neural signal changes by taking advantage of the spatial and temporal dependencies among image elements (McIntosh, Bookstein, Haxby, and Grady 1996; McIntosh and Lobaugh 2004). The adaptive boosting algorithm (AdaBoost) was proposed to classify drug-addicted subjects and healthy non drug-using controls based on observed 3D brain images (Warren and Moustafa 2022). The existence of diverse ML algorithms has raised the question of which and when an ML algorithm is better suited to extract important new information from neuroimaging data (O’Toole, Jiang, Abdi, Pénard, Dunlop, and Parent 2007; Pereira, Mitchell, and Botvinick 2009). However, the selection of the most appropriate approaches for a specific dataset and purpose poses a difficult challenge for neuroscientists with little experience in ML since it depends on the number of features used for decoding neural information as well as the effect of noisy features that are highly correlated with each other.

The curse of dimensionality of neuroimaging data can negatively affect the generalization performance of ML approaches, leading to estimation instability, model overfitting, local convergence and large estimation error. For example, the decoding abilities of algorithms that depend on specific distributions of data, such as geometric distributions of data, can be significantly influenced in the high-dimensional data space. A naïve learning approach requires the number of training data points to be an exponential function of the attribute dimension (Jain, Duin, and Mao 2000). Furthermore, the problem of high dimensionality of neuroimaging data (e.g., extremely large number of voxels) poses a number of challenges even if an approach is based on nonparametric strategies. For example, in the random forest algorithm, available features are randomly sampled to generate different subspaces of features that are used to train each decision tree in an ensemble (Kuncheva, Rodríguez, Plumpton, Linden, and Johnston 2010). Because it is typical to observe only a few features out of many that significantly contribute to the performance of these algorithms, a large number of irrelevant features appear in these subspaces. Thus, the average strength of decision trees can be diluted, thereby increasing the generalization error of the random forest algorithm. These problems also exist in neural network approaches when learning techniques are confounded by the input of high-dimensional data and the network must allocate its resources to represent many irrelevant components (Odell-Scott 1992).

Recent advancements in neuroimaging technologies have also increased the data size, namely, the total number of features to be considered has increased. The size of neuroimaging data may lead to a computational burden. However, building a pipeline for hundreds of thousands of brain regions can be very costly and time-consuming. Recently, several ML methods incorporating parallel computing environments have been developed (Xing, Ho, Xie, and Wei 2016); the implementation of a fast and efficient pipeline would be of potential application for the analysis of a large amount of neuroimaging data.

While many algorithms have been proposed, one thorough ML approach is forward variable selection (FVS). The FVS algorithm is a member of the stepwise feature selection algorithm family (Chandrashekar and Sahin 2014; Guyon and Elisseeff 2003; Kutner, Nachtsheim, Neter, and Li 2005; Weisberg 2005). It is also one of the first and most popular algorithms for causal feature selection in some fields, such as gene selection, microarray data analysis, and gene expression data analysis (Blanco, Larrañaga, Inza, and Sierra 2004; Jirapech-Umpai and Aitken 2005; Ooi and Tan 2003; Saeys, Inza, and Larranaga 2007). The powerful nature of feature decoding in the analysis of high-dimensional microbiome data has also been demonstrated (Dang and Kishino 2022). The FVS can be a powerful additional tool for neuroimaging research.

In this study, we developed a novel decoding pipeline to overcome these challenges by combining two frameworks. First, we developed an ML framework that incorporates an FVS algorithm that integrates model selection steps to detect the minimal set of features that could maximize the predictive performance. Second, the pipeline selects the best model out of a predetermined set of regression models and classifier models. This simple yet comprehensive two-stage algorithm automatically and effectively identifies important features from neuroimaging data. Moreover, because the nature of the FVS is computationally intensive and time costly, the toolbox is designed to run in a parallel environment to reduce the computational costs. Despite a significant rise in the application of ML approaches and their potential contributions to understanding brain functions, neuroimaging data are ill-posed with the high-dimensionality problem. Here, we propose a state-of-the-art and effective ML package as a solution to the high-dimensionality problem of neuroimaging data that is easy to use by neuroscientists interested in applying ML approaches to decode their neuroimaging data with little computational programming.

As a proof of concept of our approach on neuroimaging datasets, structural neuroimaging data of 1,113 subjects available in the Human Connectome Project (HCP) database were used to examine the feasibility of our proposed FVS toolbox to decode the neuroanatomical representation of biological sex and age with binary classification and multiple regression models, respectively. Structural neuroimaging data on the gray matter volume were extracted using an automated anatomical parcellation method implemented in using the CAT12 and SPM12 toolbox. For computational efficiency we reduced the dimensionality of the wholebrain voxels by extracting the gray matter volume data from 246 cortical and subcortical brain regions.

## 2. Methods and material

### 2.1. Forward variable selection (FVS) approach

The FVS approach requires an ML algorithm for feature selection and uses its performance to evaluate and determine which features are selected. The key idea behind the FVS approach is to select features that provide the largest improvement in terms of the predictive performance of the ML model, and add it to the set of selected variables in each forward iteration. The iterations stop when there is no feature improvement in the performance upon adding a feature or the maximum number of selected features has been reached. To determine whether the predictive performance of the ML model increases or decreases when a single feature is added in a greedy fashion, a number of independence tests are used.

In our approach, we used the FVS algorithm to identify a small number of features (i.e., regions of interest) to improve the performance of ML models in the subsequent step of regression or classification. At each learning step, the brain region that provides the largest increase in predictive performance and the largest reduction in MSE is selected and added to the set of selected brain regions. This process continues until there is no further performance improvement after selecting a brain region or the maximum number of selected brain regions has been reached. Model selection for brain region signature identification can also be performed using the FVS algorithm. At each forward iteration, given the selected brain regions, samples were randomly split into a training dataset comprising 70% of the samples and a test dataset comprising the remaining 30% of the samples. To select an appropriate model configuration for a specific task, fivefold cross-validation was performed on the training data to optimize the respective hyperparameters. To determine the best-performing hyperparameters for each approach, the MSE was optimized on the training dataset.

The grid search and random search strategies with cross-validation are two standard methods of hyperparameter optimization that were implemented to select the best values for the parameters of the ML model (Bisong 2019; Agrawal 2021). Based on the specific numbers of parameters of the ML model and the computational burden, the grid search with crossvalidation, in which all parameter combinations are exhaustively considered, or the random search with cross-validation, in which a given number of values are randomly selected from a parameter space, was considered for parameter optimization (Bergstra and Bengio 2012). These search strategies suffer from high-dimensional spaces but can often easily be parallelized since the hyperparameter values that the algorithm works with are usually independent of each other. For example, if ML algorithms, such as the RF algorithm and decision tree, have a large number of parameters, a random search would be an efficient option to achieve a balance between computational time and predictive accuracy. The grid search strategy is feasible for ML algorithms with a few parameters, such as the lasso or ridge models. Thus, the best ML model is specified by the selected brain regions.

The FVS algorithm is implemented in a parallel computing environment to reduce the computational burden in terms of the time cost. A number of packages provide high-performance computing solutions in Python (Palach 2014). We used the thread-based parallelism and process-based parallelism that are provided in the joblib package to separate Python worker processes to concurrently execute tasks on separate CPUs. To parallelize each of the forward iterations, the input variables were separated randomly into a number of subsets. Because of the high dimensionality of neuroimaging data, the number of these subsets (or comparisons) is usually larger than the number of processors in a single computer system. In a previous study, a computer-friendly procedure was proposed for very high-dimensional microbiome data (Dang and Kishino 2022). To introduce efficient computation, queues were created so that subsets are assigned randomly at each processor that runs the computational processes from its own privately prepared queue (Dang and Kishino 2022).

### 2.2. Regression-based ML algorithms

We explored various ML approaches to examine how a regional brain structure could contain information representing age. After the preprocessing of structural imaging data, the input (target features) included high-dimensional structural gray matter volumetric data from 1,113 samples in 246 brain regions (see Section 3 for details). We applied a variety of ML approaches to identify features and to reduce the dimension of this input with parametric regularization methods for feature selection and nonparametric methods that perform a random sampling of the available features to generate different subspaces of features to achieve a trade-off between bias and variance. In our toolbox, we selected two nonparametric methods and ten parametric methods.

For the non-parametric regression methods, the commonly used decision tree regression, random forest (RF), and Gaussian process methods were selected (Hastie, Tibshirani, Friedman, and Friedman 2009). 1) Decision tree regression is a supervised learning method that sets up a decision rule depending on the features at every interior node (Hastie *et al.* 2009). The features selected for the first partition at the root have the largest relevance. This feature selection procedure is recursively repeated for each subset at the node until further partitioning becomes impossible. The decision tree regression is typically considered to analyze MRI images (Filli, Rosskopf, Sutter, Fucentese, and Pfirrmann 2018; Kim, Kim, Shin, Yoon, Han, Koh, Roh, and Lee 2018; Naik and Patel 2014). 2) The random forest (RF) is a modification of the bagging regression that aggregates a large collection of decision trees (Breiman 2001). The main step in building an ensemble of decision trees is to perform random sampling of the available features to generate different subspaces of features at each node of each unpruned decision tree. Using this strategy, better estimation performances can be obtained compared with using single decision tree because each tree estimator has low bias but high variance, whereas a bias-variance trade-off is achieved by the bagging process of RF. The random forest (RF) algorithm has become a standard data analysis tool in multiple areas such as bioinformatics (Boulesteix, Janitza, Kruppa, and König 2012; Zhang and Ma 2012) and neuroimaging analysis (Eshaghi, Wottschel, Cortese, Calabrese, Sahraian, Thompson, Alexander, and Ciccarelli 2016; Mitra, Bourgeat, Fripp, Ghose, Rose, Salvado, Connelly, Campbell, Palmer, Sharma *et al.* 2014). (3) The gaussian process (GP) is a nonparametric model that is a natural generalization of a multivariate Gaussian distribution to a Gaussian distribution over a specific family of functions such as kernel functions. In GP regression, a prior distribution is proposed directly over the nonlinear function space, rather than specifying a parametric family of nonlinear functions. Different kernels can be used to express different structures observed in the data. Thus, the GP has a large degree of flexibility in capturing the underlying signals without imposing strong modeling assumptions. This property makes the GP an attractive model for analyzing genetic data (Chu, Ghahramani, Falciani, and Wild 2005) as well as MRI data (Wassermann, Bloy, Kanterakis, Verma, and Deriche 2010).

We selected eight parametric ML algorithms including ridge regression, least absolute shrinkage and selection operator (Lasso) regression, kernel ridge regression, multitask Lasso regression, least angle regression (Lar), LassoLar regression, elastic net regression, and regularized linear models with stochastic gradient descent (SGD). 1) Ridge regression is a linear least squares method that uses L2 regularization or weight decay to control the relative importance of features (Hastie *et al.* 2009). L2 regularization encourages weight values to decay toward zero. Thus, ridge regression can be used to overcome the disadvantages of the ordinary least square method, i.e., that the variance in the estimate of the linear transform may be large because the number of features is significantly larger than the number of samples. 2) Least absolute shrinkage and selection operator (Lasso) regression, which is another type of linear regression, uses L1 regularization and can eliminate a number of coefficients from the model by adding a penalty equal to the absolute value of their magnitude (Hastie *et al.* 2009). 3) Kernel ridge regression is an extension of ridge regression that is used when the number of dimensions can be much larger, or even infinitely larger, than the number of samples (Schölkopf, Luo, and Vovk 2013). The main idea is to propose the kernel trick to convert the original data space into the fancy feature space that can significantly reduce the computational burden of learning processes. 4) Multitask Lasso regression generalizes the Lasso to the multitask setting by replacing the L1-norm regularization term with the supnorm regularization sum (Hastie *et al.* 2009). 5) The least angle regression (Lar) model is the modification of the Lasso and the forward stagewise linear regression approaches, where the number of features is significantly greater than the number of samples (Efron, Hastie, Johnstone, and Tibshirani 2004). At each iteration, Lars selects the feature most correlated with the target. If multiple features have a similar correlation, the direction equiangular between the features is moved forward. 6) The Lasso model fit with least angle regression (LassoLar) is the combination of Lar and Lasso and is implemented to improve the variable selection (Efron *et al.* 2004). 7) The elastic net model is the generalization of ridge regression and lasso. This model proposes the elastic net penalty, which controls the balance between L1 and L2 regularization of the coefficients (Hastie *et al.* 2009; Zou and Hastie 2005). Thus, elastic net can be used to perform feature selection in a high-dimensional space. and 8) Regularized linear models with stochastic gradient descent (SGD) learning are an extension of the ridge, Lasso and elastic net approaches with a large number of training samples implemented with a plain stochastic gradient descent learning routine (Hastie *et al.* 2009). We provide the specific explanations for each ML algorithm in the Supplementary Material.

For each ML algorithm, the parameter optimization steps were developed. Specifically, in the cases of the ridge, Lasso, multitask Lasso, Lar, and LassoLar algorithms, the complexity of the parameters that control the amount of shrinkage was optimized. The elastic net model includes an additional parameter that controls a combination of L1 and L2 penalties separately. The main parameters of the regularized linear models with SGD learning were loss functions, penalty options, and the learning rate schedule, whereas those of the kernel ridge regression model were kernel options that include linear, Laplacian, Gaussian, and sigmoid kernels, the regularization parameter, and the kernel coefficient, and those of the decision tree model were the maximum depth of the tree, the minimum number of samples required to split an internal node, the minimum number of the samples required to be at a leaf node, the function to measure the quality of a split, the strategy used to choose the split at each node, and the number of features to consider when looking for the best split. The parameters of the random forest regressor were the same as those of the decision tree regressor, except that the number of trees was an additional parameter.

As a criterion to compare the performance of regression, the mean squared error (MSE), mean absolute error (MAE), and Spearman correlation coefficients that were calculated between the predicted and the true values were calculated (Hastie *et al.* 2009). In the field of statistics, the Akaike or Bayesian information criteria (also known as AIC or BIC, respectively) are widely used indices to quantify the fit of a model (Burnham and Anderson 2004); however, these information criteria methods do not apply for nonparametric regression models (e.g., decision tree, random forest). Thus, we selected the MSE and MAE, which are applicable across all models.

### 2.3. Classification-based ML algorithms

We also extended the application of our approach to study the binary or multiclass classification. We examined how brain structure could contain information representing sex (male or female). The automated classification algorithms include nonparametric methods, such as decision tree, random forest, gradient boosting method, extreme gradient boosting, and extremely randomized trees, which have the capability of regression and classification. 1) Decision trees are commonly utilized classification models in various fields, such as machine learning and data mining (Gavankar and Sawarkar 2017). Decision trees include a number of tests or attribute nodes linked to subtrees and decision nodes labeled with a class, i.e., a decision. A sample is classified by starting at the root node of the tree. Each node represents features in a group to be classified and each subset defines a value that can be taken by the node (Hastie *et al.* 2009). The entropy, Gini index, and information gain are the standard measures of a dataset’s impurity or randomness in decision tree classification. 2) Random forest classification is one of the most popular ensemble methods that can be used to avoid the tendency of simple decision trees to overfit (Breiman 2001). Similar to regression, random forest classification proposes a slightly randomized training process to independently build multiple decision trees. The randomization processes include using only a random subset of the whole training dataset to build each tree and using a random subset of the features or a random splitting point when considering an optimal split. 3) The gradient boosting method is an ensemble approach that uses the boosting technique to combine a sequence of weak decision trees (Friedman 2001). Each tree in the gradient boosting fits to the residuals from the previous tree. Thus, the errors of the previous tree are minimized, and the overall accuracy and robustness of the model are considerably improved. 4) Extreme gradient boosting is an efficient and scalable implementation of the gradient boosting framework for sparse data with billions of examples (Chen and Guestrin 2016). 5) Extremely randomized trees are another approach to improve the performance of decision trees by generating diverse ensembles (Geurts, Ernst, and Wehenkel 2006). The main idea of this approach is to inject randomness into the training process by selecting the best splitting attribute from a random subset of features. However, in contrast to random forest, the bootstrap instances procedures are implemented by extremely randomized trees. We provide specific explanations for each ML algorithm in the Supplementary Material.

Because of the computational burdens of nonparametric approaches, random search strategies with cross-validation are implemented for the parameter optimization steps. The parameters of decision tree classification, such as the maximum depth of the tree and the minimum number of samples required to split an internal node, are similar to those of regression. The Gini index and entropy are used to measure the quality of a split in classification. Moreover, a large number of parameters of random forest, gradient boosting, extreme gradient boosting, and extremely randomized trees are similar to the parameters of decision tree classification. However, there are several special parameters that can significantly influence performance. Specifically, the number of trees is the most important parameter of random forest classification. The necessary parameters of gradient boosting classification include the loss function to binomial and multinomial deviance, the function to measure the quality of a split, the function to measure the quality of a split, and the number of boosting stages. The parameters of extreme gradient boosting are similar to those of gradient boosting. Because it has an option for the number of parallel trees constructed during each iteration, its computational speed is faster. An important parameter of extremely randomized trees is the number of trees in the forest, and the bootstrapping technique is not used to build each tree.

Simple parametric models, such as logistic regression and naïve Bayes, were also included in the classification algorithms. 1) Logistic regression is a standard method for building prediction models for classification. Due to the high-dimensional problems of multiple areas (Bühlmann and Van De Geer 2011), ridge and Lasso penalties are added to penalized logistic modeling for the feature selection step. This method has been applied for the analysis of genetic datasets to select a subset of genes that can provide more accurate diagnostic methods (Liao and Chin 2007; Wu, Chen, Hastie, Sobel, and Lange 2009). 2) Naïve Bayes is a classification model that refers to the construction of a Bayesian probabilistic model to assign a posterior class probability to each sample (McCallum, Nigam *et al.* 1998). The important assumption of this method is that the features constituting the sample are conditionally independent given the class. The naïve Bayes method is fast, easy to implement and relatively effective for the classification of biological datasets (Yousef, Jung, Kossenkov, Showe, and Showe 2007). The grid search strategy with cross-validation is implemented for the parameter optimization steps. The parameter in logistic classification is an elastic net mixing parameter to control the combination of the L1 and L2 regularization. The naïve Bayes classification parameter is an additive (Laplace/Lidstone) smoothing parameter.

To evaluate the decoding performance, three main criteria were compared across tested models: ‘precision’ is defined as the number of true positives over the number of true positives plus the number of false-positives, ‘recall’ is defined as the number of true positives over the number of true positives plus the number of false-negatives, and the ‘F1 score’ is defined as the harmonic mean of precision and recall. 1 depicts the steps for searching for important features using FVS, the application of each ML model and how these computations are appropriately decomposed in a parallel computation manner.

### 2.4. Neuroimaging data samples

To examine the feasibility of the proposed pipeline in neuroimaging, we acquired high-resolution structural MRI scans of a large number of healthy subjects from the Human Connectome Project (HCP). This dataset includes 1,113 samples. The structural images were segmented into gray matter, white matter, and cerebrospinal fluid and normalized (1 x 1 x 1 voxel size) into a template space using standard parameters implemented in the Computational Neuroanatomy Toolbox (CAT12). During the segmentation process, CAT12 implemented an automated parcellation of the gray matter to extract the gray matter volume in native space from 246 cortical and subcortical brain regions according to neuroanatomical landmarks based on the Brainnetome Atlas (https://atlas.brainnetome.org/bnatlas.html) (Fan, Li, Zhuo, Zhang, Wang, Chen, Yang, Chu, Xie, Laird *et al.* 2016). CAT12 was also used to estimate individual values of the total intracranial volume (TIV), which was included as a covariate of no interest for the classification and regression models. Notably, the pipeline technically works for the whole-brain voxel-based dataset; however, these segmented data were used for simplicity.

Here, we provide a use case example to identify the best model to predict the target variable. More specifically, the gray matter volume data from 246 Brainnetome regions were selected as target features to predict the age and sex of participants using regression and classification models, respectively.

## 3. Package structure

Our framework includes two core modules: automatic ML and FVS algorithms for regression and classification. First, ML models and FVS algorithms were implemented using the Python programming language to optimize the parallel computations that could significantly reduce the computation time. The scikit-learn library in Python was used to implement core computational techniques for the random forest classifier. Our Python package to implement the proposed approach is available on Github (https://github.com/tungtokyo1108/FVS_decoder).

In general, each user creates a short script of regression or classification that contains (1) automatic ML algorithms for the input dataset and (2) the FVS algorithm combined with the best ML algorithm in step (1). For example, the script for regression after controlling the effects of variables such as the total intracranial volume (TIV) is short.

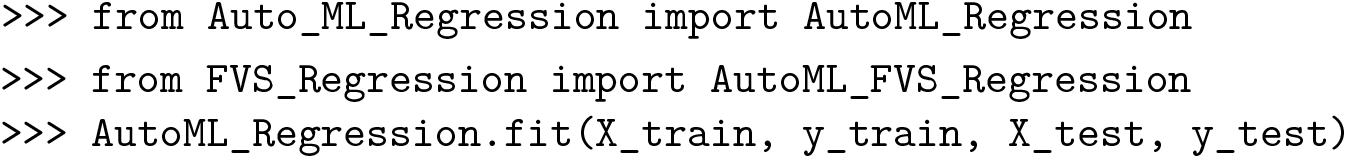

This function runs 11 ML regression algorithms to select the best algorithm for the input dataset. The output of this function is a table that shows the rank of performances of 11 ML regression algorithms based on their performance.

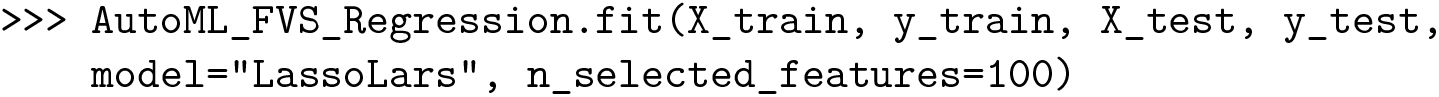

After selecting the best ML algorithm, the user implements the function that runs the FVS algorithm to identify an important group of ROIs. For example, the LassoLars algorithm is the best model with the smallest value of MSE in our dataset. Thus, we want to combine the LassoLars model with the FVS algorithm and the maximum number of features, that we want to select is 100. We define model as “LassoLars”, n_selected_features=100. The details of the parameters and outputs of all functions in our package are provided in the README.md file on Github. Table 1 shows the main functions of our package. These functions are then applied to real data in Section 4.

**Table 1:**
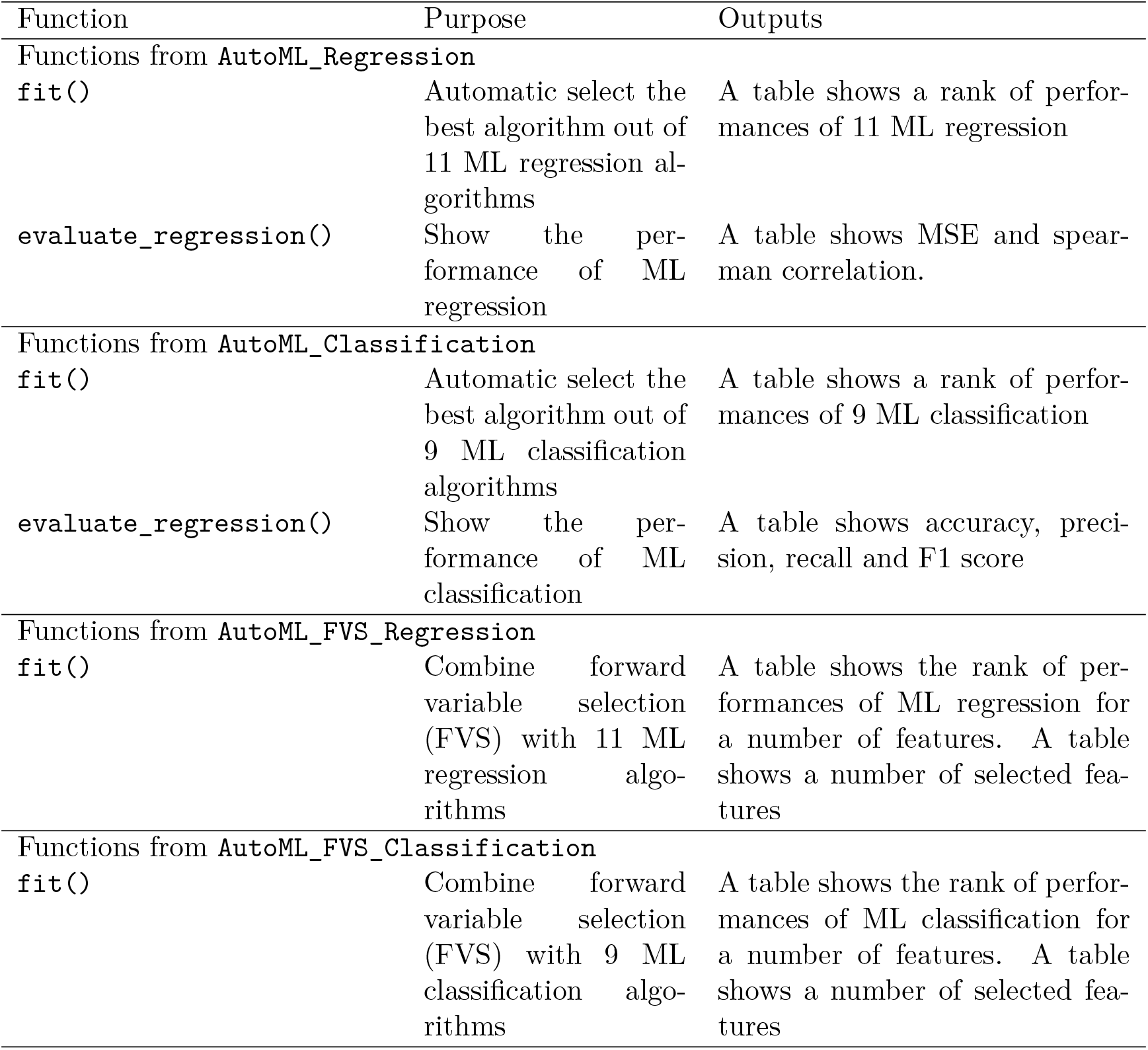
An overview of the main functions in FVSdecoder package.

| Function | Purpose | Outputs |
| --- | --- | --- |
| Functions from AutoML_Regression |  |  |
| `fit()` | Automatic select the best algorithm out of 11 ML regression algorithms | A table shows a rank of performances of 11 ML regression |
| `evaluate_regression()` | Show the performance of ML regression | A table shows MSE and spearman correlation. |
| Functions from AutoML_Classification |  |  |
| `fit()` | Automatic select the best algorithm out of 9 ML classification algorithms | A table shows a rank of performances of 9 ML classification |
| `evaluate_regression()` | Show the performance of ML classification | A table shows accuracy, precision, recall and F1 score |
| Functions from AutoML_FVS_Regression |  |  |
| `fit()` | Combine forward variable selection (FVS) with 11 ML regression algorithms | A table shows the rank of performances of ML regression for a number of features. A table shows a number of selected features |
| Functions from AutoML_FVS_Classification |  |  |
| `fit()` | Combine forward variable selection (FVS) with 9 ML classification algorithms | A table shows the rank of performances of ML classification for a number of features. A table shows a number of selected features |

## 4. Illustrations

### 4.1. Improved accuracy for prediction of age from MRI data

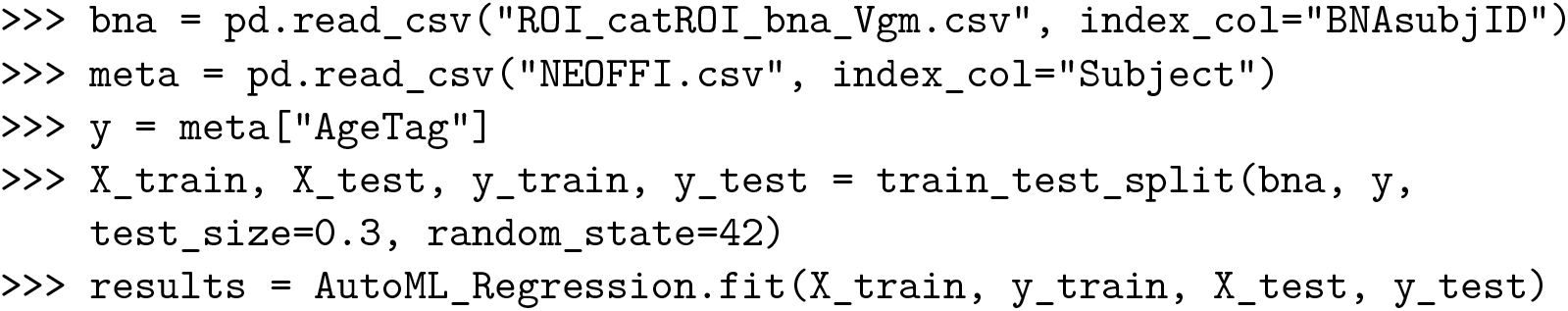

Table 2 summarizes the MSE values for each ML algorithm with and without the FVS algorithm to predict the age of healthy individuals. For the comparisons without FVS, the best performance and the smallest MSE (MSE = 0.3686) were obtained using the LassoLars regression model.

**Table 2:** Accuracies of the ML models as assessed by MSE to predict age. FVS: forward variable selection algorithm; MSE: mean squared error.

| Model | MSE without FVS | MSE with FVS |
| --- | --- | --- |
| LassoLar | 0.4541 | 0.3686 |
| Random Forest | 0.4807 | 0.3722 |
| Gaussian Process | 0.4855 | 0.3895 |
| Ridge | 0.4900 | 0.3928 |
| Elastic net | 0.4909 | 0.4011 |
| Lars | 0.4988 | 0.4171 |
| Lasso | 0.5034 | 0.4265 |
| Kernel Ridge | 0.5061 | 0.4402 |
| Multitask Lasso | 0.5341 | 0.4578 |
| Decision Tree | 0.5669 | 0.4669 |
| Stochastic Gradient Descent | 0.5687 | 0.5000 |

Across the 11 models, the use of FVS methods significantly improved the performance compared with not using the FVS methods (Figure 2). In addition, the random forest and Gaussian process algorithms showed a comparable accuracy with the LassoLars regression model. However, the computational costs of the random forest (CPU time = 2 minutes without FVS) and Gaussian process (CPU time = 80 seconds without FVS) methods were more excessive because the parameters were more complex than those of LassoLars regression (CPU time = 20 seconds without FVS). The other approaches, such as ridge, elastic net and Lars regression, had faster computations, but good performance could not be obtained using these approaches. Therefore, we focused the LassoLars regression model for the next step of analysis on the HCP dataset. All parallel computations were run on an Apple M1 Max with 10 CPU cores.

**Figure 1:**
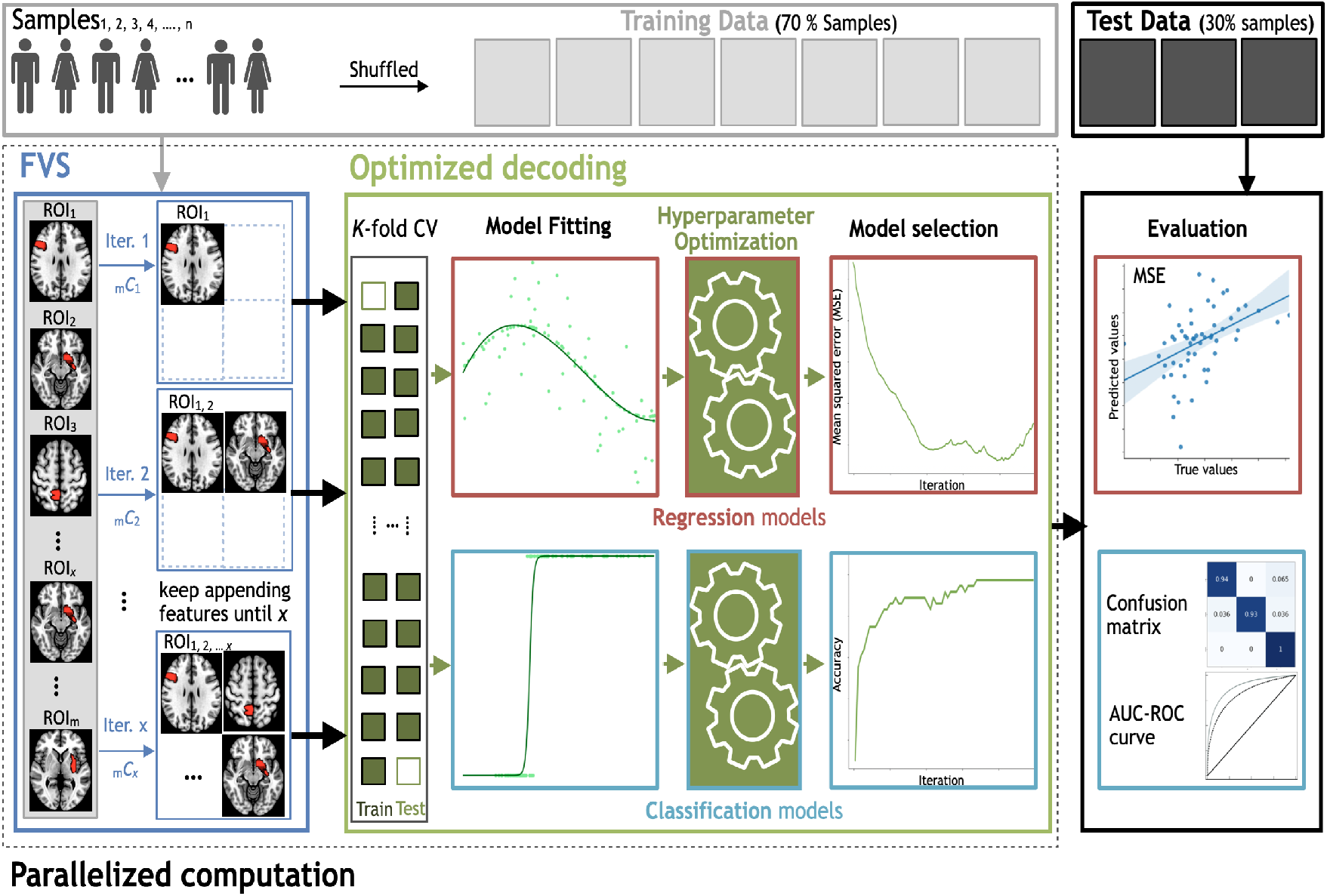
Workflow schematics of the automatic ML toolbox coupled with the forward variable selection (FVS) algorithm. All features, i.e., the gray matter volume data from each region of interest (ROI), undergo the FVS step, followed by either regression-based or classification-based ML with K-fold cross-validation (CV). The random search and grid search strategies with cross-validation were adopted to optimize the hyperparameters of the ML algorithms at each iteration of the FVS algorithm. The final outcomes were evaluated based on the MSE and MAE for regression-based models and the AUC and confusion matrix for classification-based model. n is the number of samples, m is the total number of ROIs (246 ROIs in this study) and x is the number of ROIs that the user wants to select.

**Figure 2:**
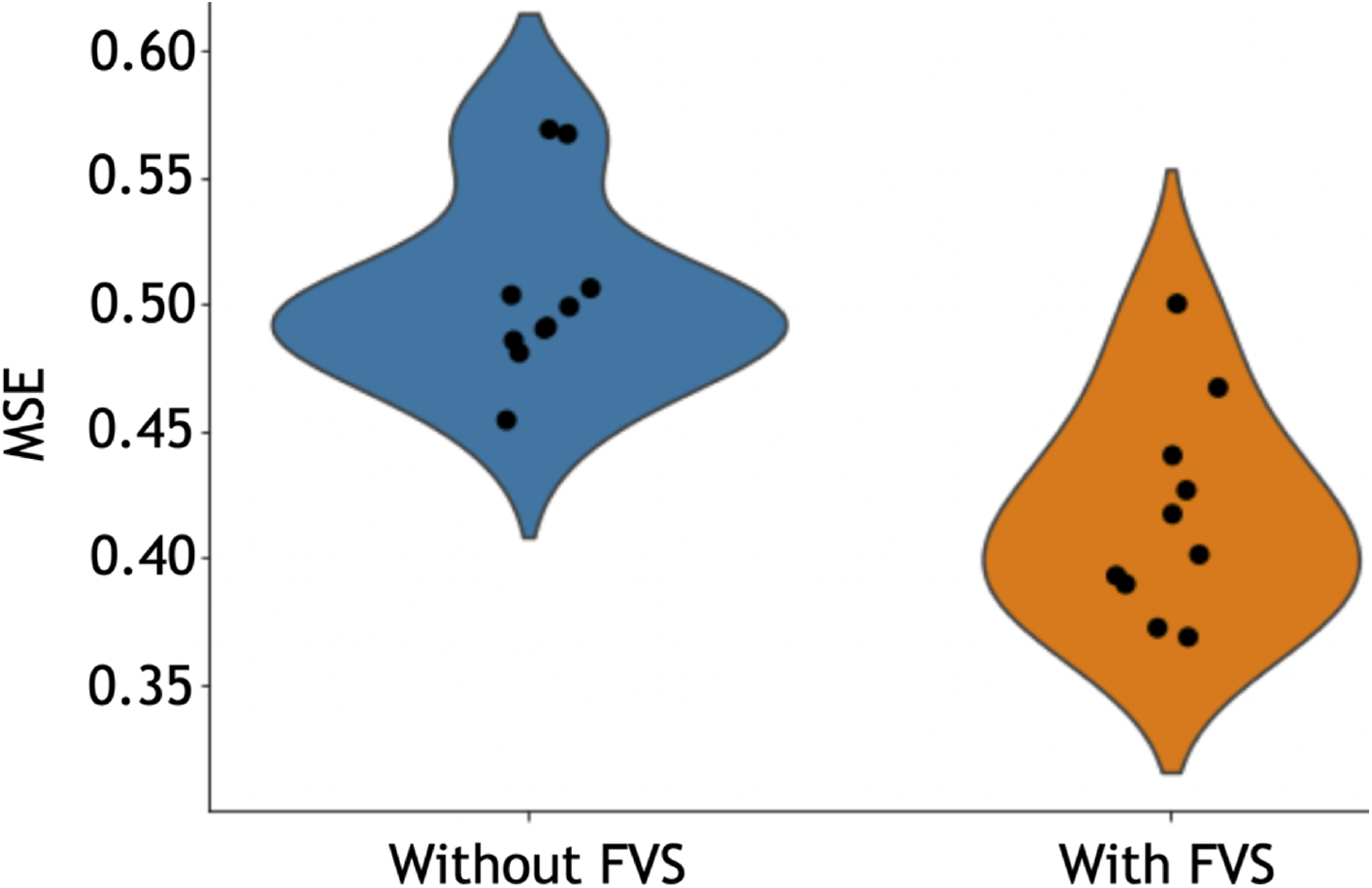
Performance comparison of 11 regression models with and without forward variable selection (FVS) to predict age, controlling for TIV. Left (blue): 11 regression models without the FVS algorithm. Right (orange): 11 regression models on a subset of brain regions selected with the FVS algorithm.

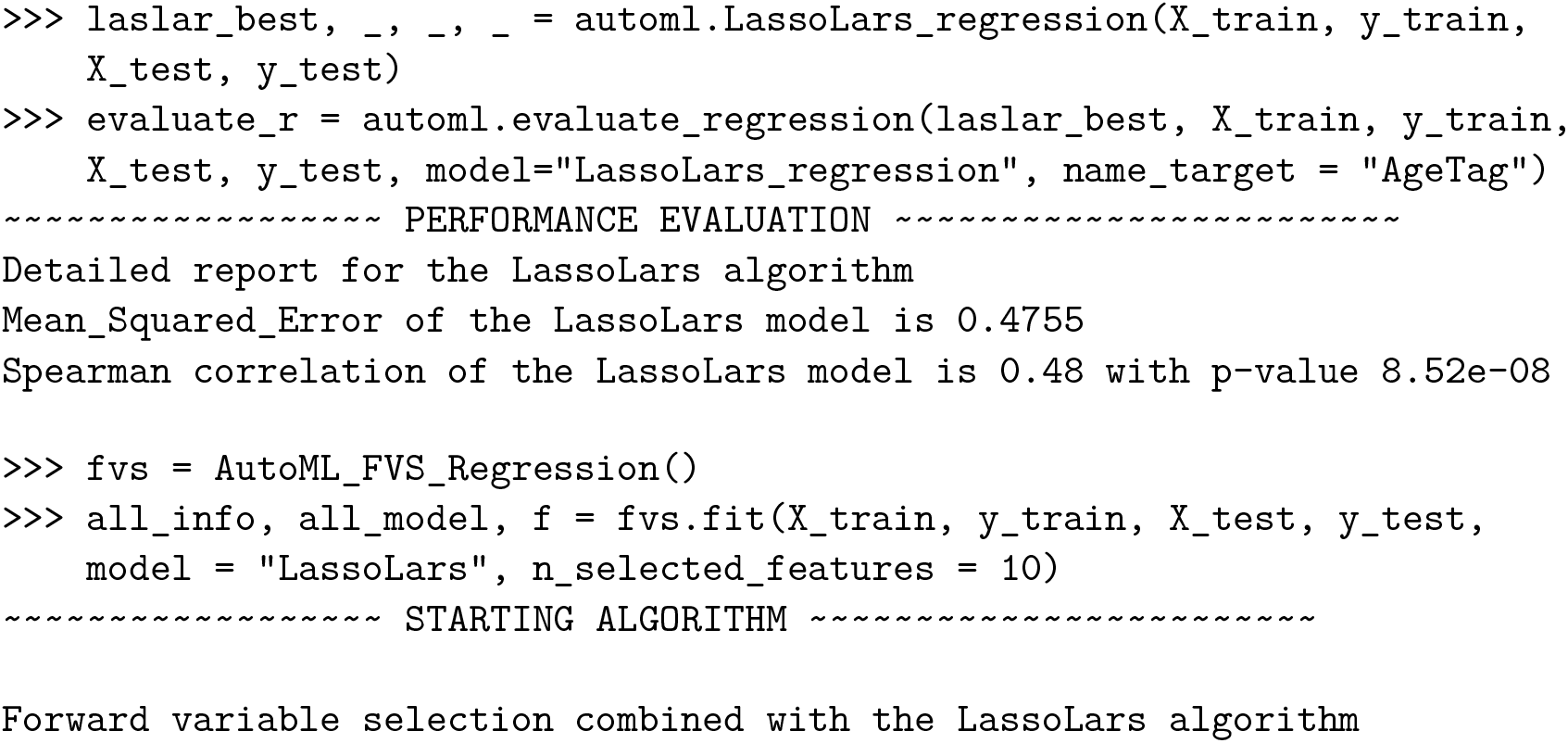

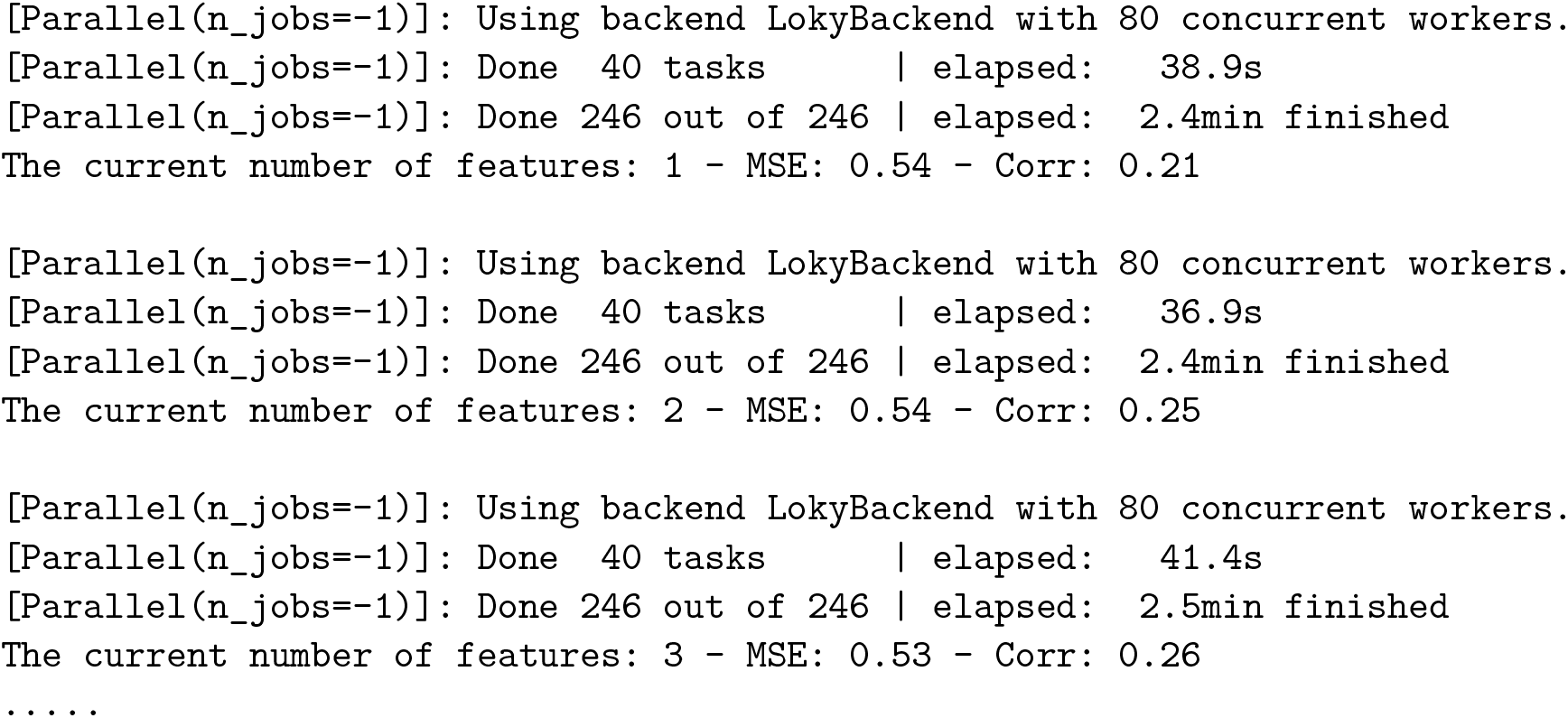

Among all model comparisons, 54 out of 246 brain regions were identified with a Spearman correlation coefficient of .63 (p < .0001, Figure 3) using the FVS-supported LassoLar regression model. Figure 4 shows selected brain regions identified by the FVS-supported LassoLars model. These findings were consistent with previous reports of a significant association of brain regions with age. For example, the thalamus plays a critical role in the coordination of information flow in the brain, mediating communication and integrating many processes, including memory, attention, and perception. Thus, age-related cognitive capability could be associated with micro- and macrostructural alterations in regions of the thalamus. A number of previous studies have shown that increasing age significantly influences the changes in thalamic shape and in the volume (Good, Johnsrude, Ashburner, Henson, Friston, and Frackowiak 2001; Hutton, Draganski, Ashburner, and Weiskopf 2009).

**Figure 3:**
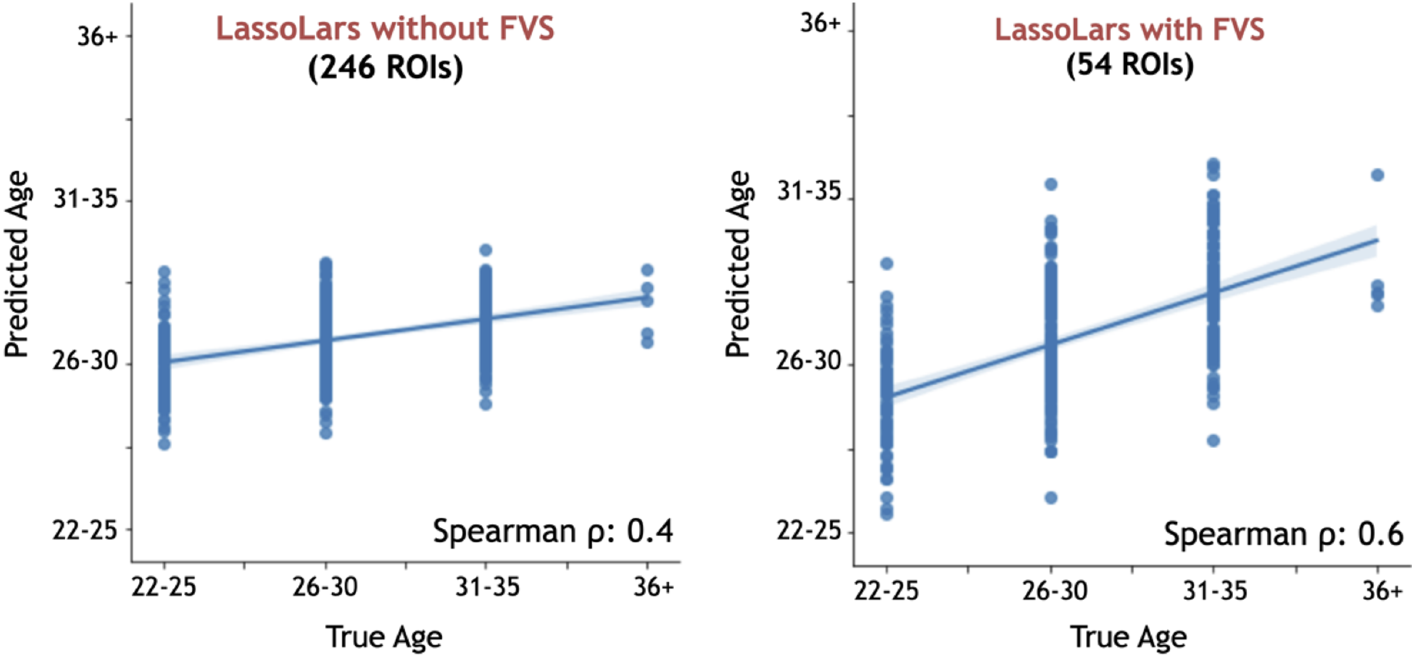
Performance comparison of LassoLar regression with and without the forward variable selection (FVS) algorithm to predict age, controlling for the effects of TIV. Left panel: LassoLar regression analysis with all of brain regions (MSE = 0.45, Spearman *ρ* = .44, p = .064). Right panel: LassoLar regression on a subset of brain regions selected with the FVS algorithm (MSE = 0.36, Spearman *ρ* = .63, p < .0001). Predicted age data are plotted as a function of the true score. The blue lines and blue shades represent a liner regression line with confidence interval.

**Figure 4:**
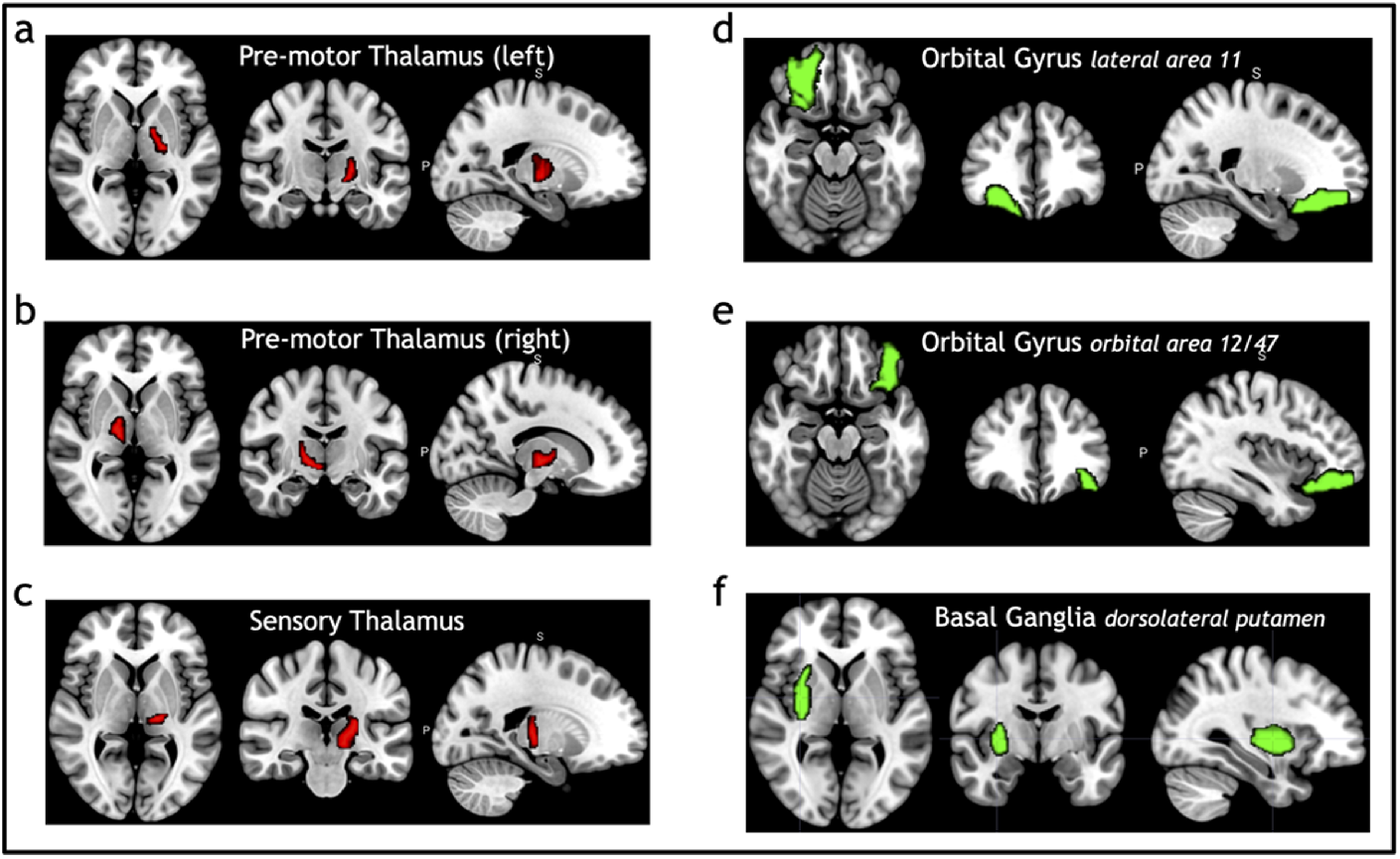
Selected brain regions significantly associated with age. The red color denotes a positive correlation with age; the green color denotes a negative correlation with age. a: premotor thalamus (left), b: premotor thalamus (right), c: sensory thalamus (left), d: orbital gyrus lateral area 11, e: orbital gyrus orbital area 12/47, f: basal ganglia dorsolateral putamen.

### 4.2. Improved accuracy for fMRI data from classification study

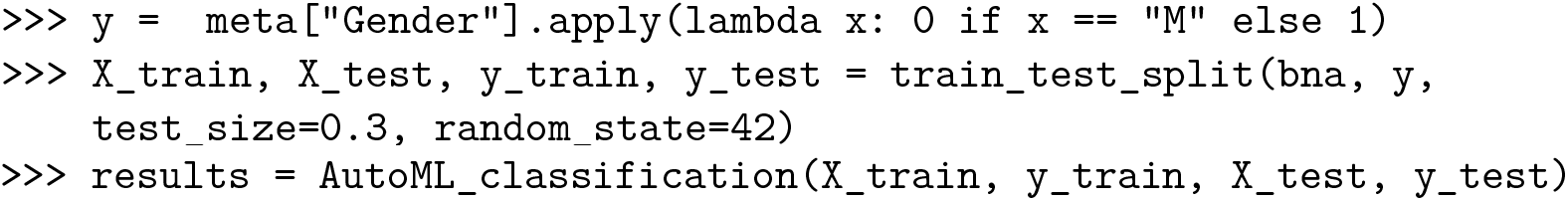

Table 3 summarizes the accuracies for each ML algorithm with and without the FVS algorithm that classifies the male and female groups. The best performance among the comparisons without FVS, with the highest accuracy of 75.44%, was obtained using the random forest classifier.

**Table 3:**
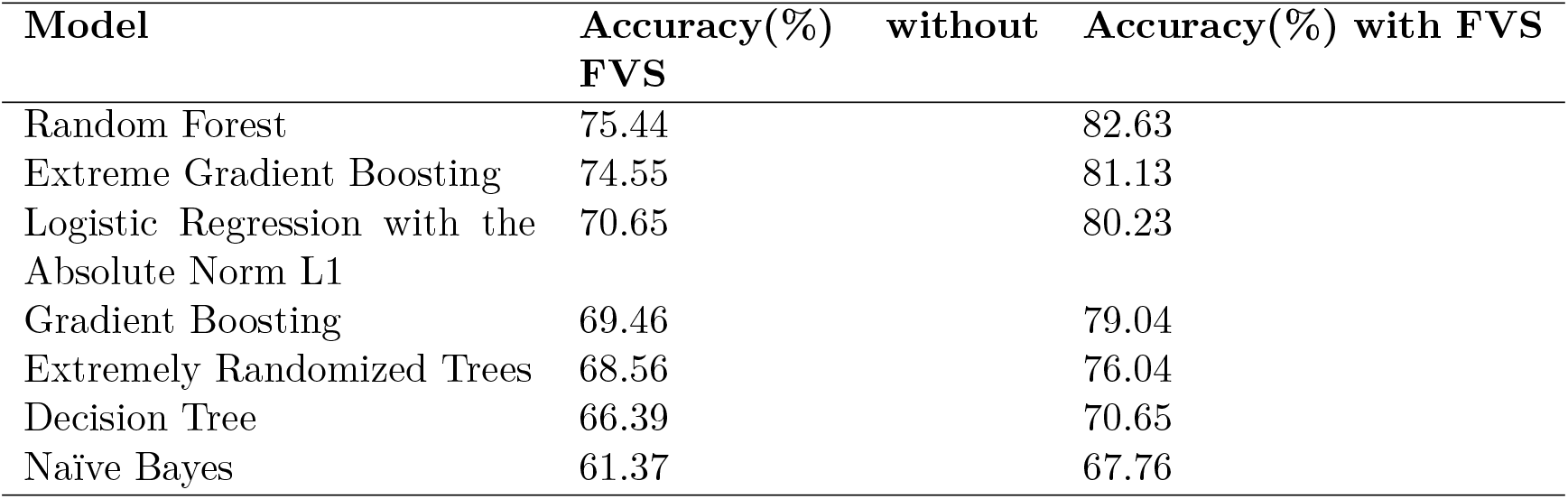
Accuracies of the ML models used to classify the male and female groups. FVS: forward variable selection algorithm.

| Model | Accuracy(%)<br>without FVS | Accuracy(%) with FVS |
| --- | --- | --- |
| Random Forest | 75.44 | 82.63 |
| Extreme Gradient Boosting | 74.55 | 81.13 |
| Logistic Regression with the Absolute Norm L1 | 70.65 | 80.23 |
| Gradient Boosting | 69.46 | 79.04 |
| Extremely Randomized Trees | 68.56 | 76.04 |
| Decision Tree | 66.39 | 70.65 |
| Naïve Bayes | 61.37 | 67.76 |

Figure 5 shows that across the seven classifiers, the use of FVS significantly improved the performance compared with not using FVS. In addition, the gradient boosting algorithm in Table 3 showed the comparable accuracy (74.55%) with the random forest classifier. However, the computational costs of the extreme gradient boosting (CPU time = 4.5 minutes without FVS) method were more expensive because the parameters were more complex than those of the random forest classifier (CPU time = 2 minutes without FVS). Additionally, logistic regression with the absolute norm L1 achieved a comparable performance to the extreme gradient boosting classifier with high accuracy (70.65%), while the computational burden was reduced considerably (CPU time = 19 seconds without FVS). Conversely, the extremely randomized trees and naïve Bayes algorithms had poor performances with low accuracy values (68.56% and 61.37%).

**Figure 5:**
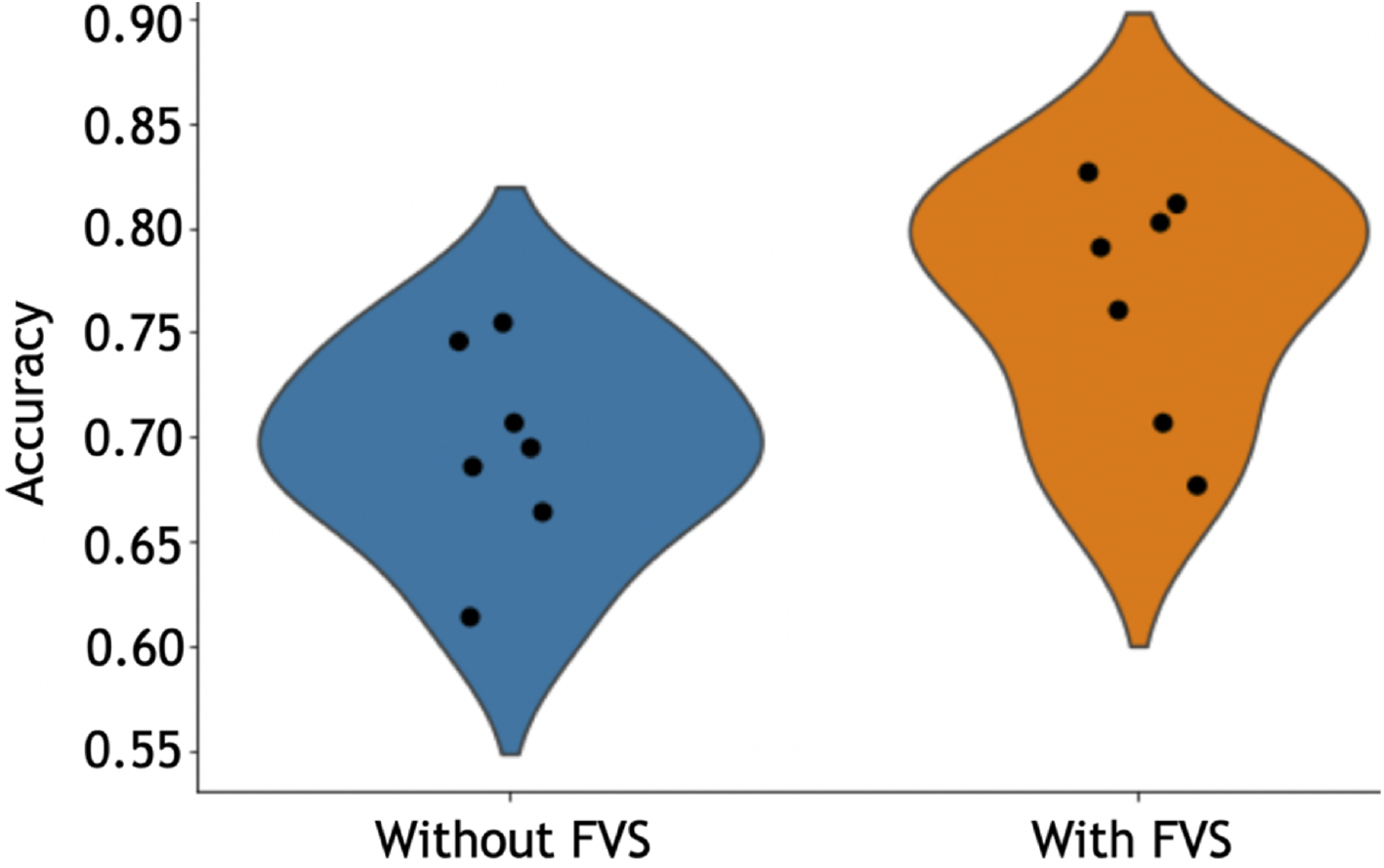
Performance comparison of 7 classification models with and without the forward variable selection (FVS) algorithm to classify male and female groups, controlling for the effects of TIV. Left hand (blue): 7 regression models without the FVS algorithm. Right hand (orange): 7 regression models on a subset of brain regions selected by the FVS algorithm.

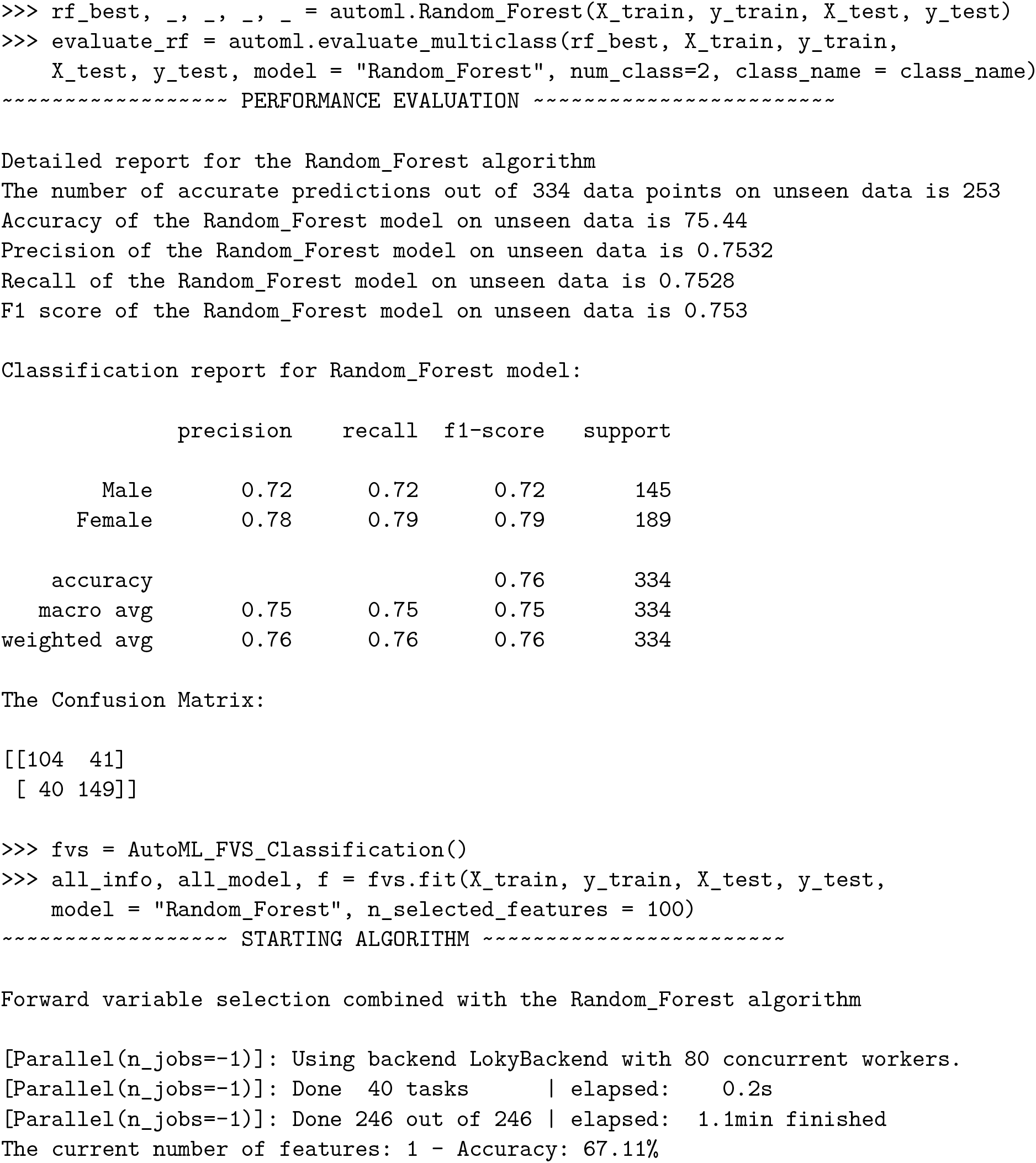

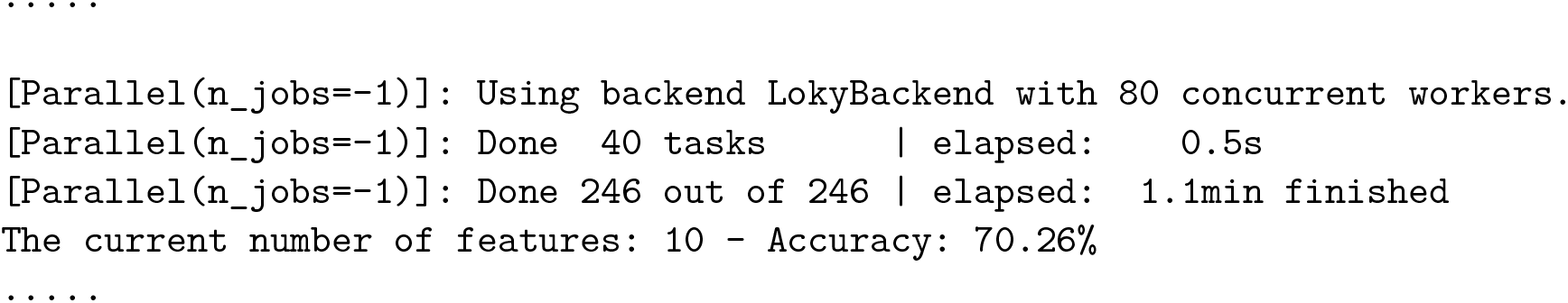

Among all model comparisons, Figure 6 shows that 87 out of 246 brain regions were identified and the accuracy improved to 82.63% using the FVS-supported random forest classifier. Females were identified with an accuracy of 87%, and males were identified with an accuracy of 77%. Figure 7 shows that the selected brain regions, such as the thalamus, inferior frontal gyrus, precuneus, and basal ganglia, were mapped on the Brainnetome Atlas. These brain regions were identified by our model and were consistent with previous reports. For example, a number of studies showed that females had significantly greater volumes in the inferior frontal gyrus, thalamus and precuneus. Conversely, males had significantly greater volumes in the basal ganglia and medio ventral occipital.

**Figure 6:**
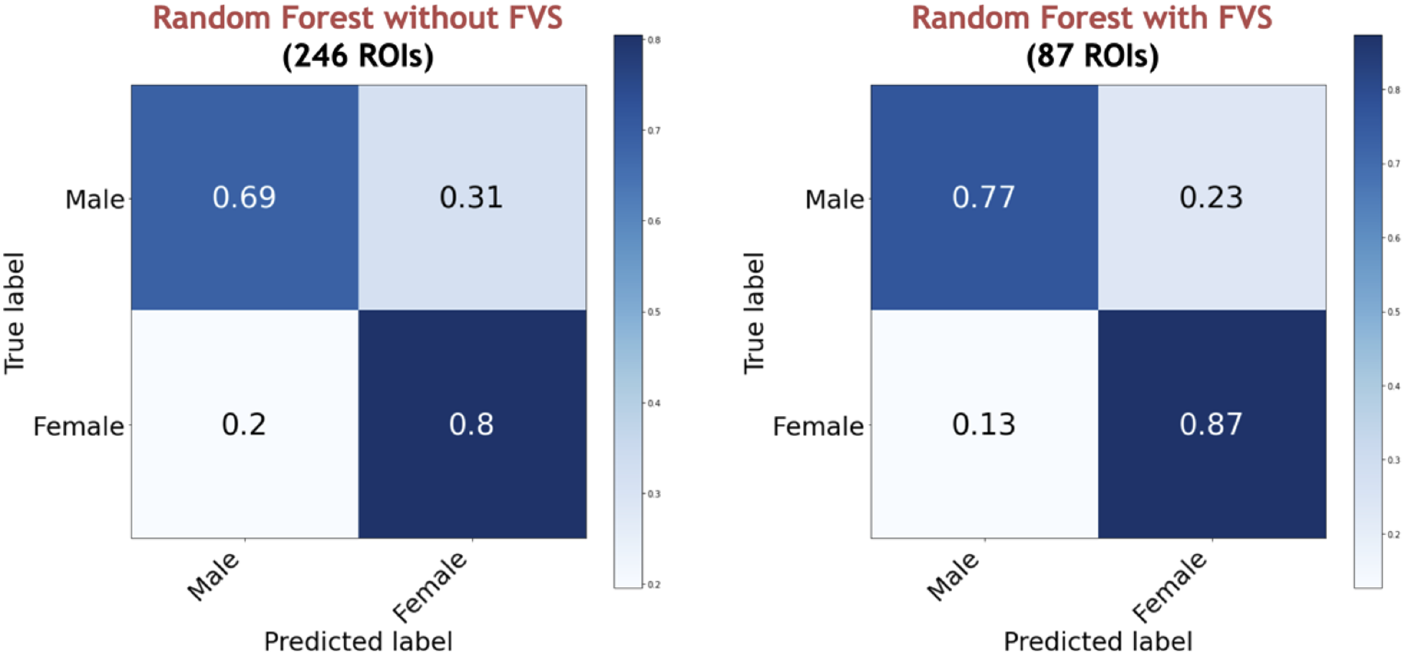
Performance comparison of the random forest classifier with and without the forward variable selection (FVS) algorithm to classify two groups, controlling for the effects of TIV. Left panel: random forest classifier analysis with all of brain regions. Right panel: random forest classifier on a subset of brain regions selected by the FVS algorithm.

**Figure 7:**
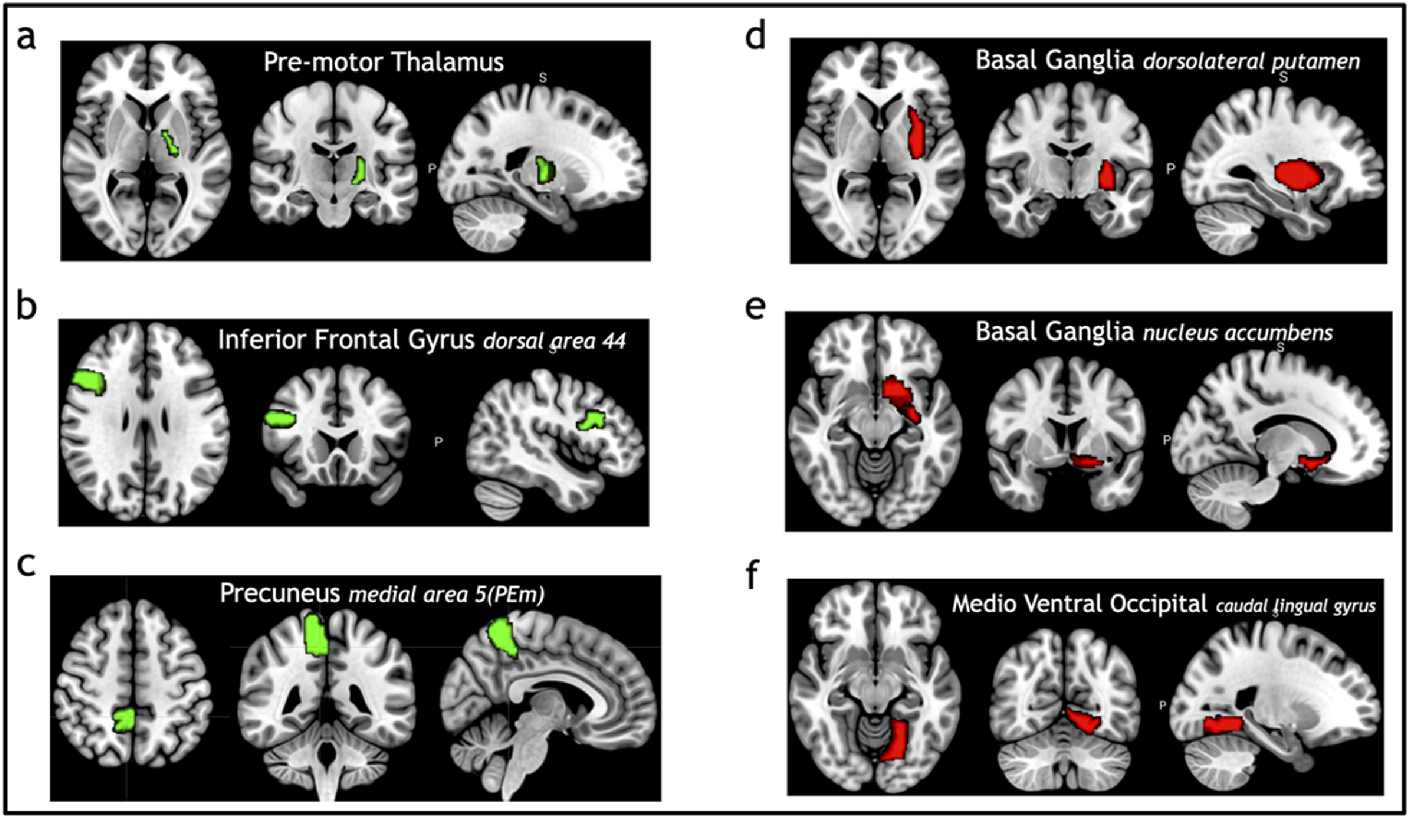
Selected brain regions identified as predictors of the sex categories (male and female). The red color denotes male predicting volume > female predicting volume; the green color denotes male predicting volume < female predicting volume. a: premotor thalamus, b: inferior frontal gyrus dorsal area 44, c: Precuneus medial area 5(PEm), d: Basal ganglia dorsolateral putamen, e: Basal ganglia nucleus accumbens, f: Medio ventral occipital caudal lingual gyrus.

## 5. Summary and discussion

In this study, a parallelized FVS toolbox is developed to provide optimized decoding of neuroimaging data samples. Our toolbox can be used to propose the best ML model for user’s input data and identify a small group of important features that significantly improve the performance of the ML algorithm. We have demonstrated that the toolbox is feasible for region of interest (ROI) data without revising the model types (parametric or nonparametric) and parameter settings, suggesting that this toolbox is generalizable and could potentially be used to train multiple types of neuroimaging data without modification. Given previous use cases of the ML approach that have been established in genetic studies using the FVS algorithm (Dang and Kishino 2022), we have extended the FVS algorithm, and the toolbox has been created for neuroimaging studies. To examine the feasibility of our ML pipelines, sample neuroimaging data were acquired from the HCP database. As case samples, we compared the accuracies (predictability) of the classical ML algorithms with and without FVS.

We tested the performances of several ML models by analyzing large structural MRI datasets with a large number of variables (246 brain regions). An easy-to-use computational package may help novel data scientists in neuroimaging research and advance the research by identifying accurate features relevant to questions of interest.

### 5.1. Comparison against existing methods

The proposed approach presents the following advantages compared with the previous methods. First, neuroscientists could avoid decision uncertainties when considering or choosing the most appropriate model for their own datasets. In our proposed approach, users only provide the input data and decide whether to run the proposed ML pipeline for either classification or regression based on their purpose of analysis. The automatic algorithm will rank the ML algorithms and recommend the best algorithm for the user’s dataset. In this study, the results showed that random forest was the most accurate algorithm for classification. Random forest is an ML algorithm that is based on combining multiple decision trees by random selection of samples. Therefore, random forest overcomes the problem of overfitting decision trees that can result in a better fitting of the model (Ghose, Mitra, Oliver, Marti, Lladó, Freixenet, Vilanova, Sidibé, and Meriaudeau 2012; Mitra *et al.* 2014; Sarica, Cerasa, and Quattrone 2017; Zhu, Du, Kerich, Lohoff, and Momenan 2018). In the regression task, the best performance for predicting the age of healthy individuals was obtained using the LassoLar method. The performance of random forest was ranked second (Dimitriadis, Liparas, Tsolaki, Initiative *et al.* 2018; Jog, Carass, Roy, Pham, and Prince 2017; Smith, Ganesh, and Liu 2013). In second step, the FVS algorithm was used to select a feature (e.g., ROI) that improves the accuracies of ML classification algorithms or reduces the MSEs of ML regression algorithms at each iteration. This procedure was stopped if the performance of the ML algorithm reached a maximization. The FVS algorithm attempts to identify a minimal core set of brain regions that can provide insights into brain functions. The results showed that the performances of all ML algorithms in classification and regression were significantly improved after applying the FVS algorithm. For example, the FVS algorithms identified 87 ROI features that improved the accuracy of the random forest classifier from 75.44% to 82.63%.

### 5.2. Limitations

Although it was apparent that the FVS algorithm robustly and significantly improved the accuracy for both classification and regression models, the downside of this approach is the computation time to apply nearly all possible pairs to consider all features (up to the specified number of pairs specified by the user). To compensate for the issue of time, our toolbox has implemented parallel computing pipelines to effectively minimize the computational time.

In addition, there are some important limitations to the generalizability of the findings in this study. While the performances of the ML algorithm are significantly improved by the FVS algorithm, computational burden of the FVS algorithm is still a difficult challenge for personal computers. Even if we apply the parallel computational techniques to overcome large-scale problems in the FVS algorithm, a high-performance computer is necessary to efficiently run our proposed tool. Although the computational speed could be improved, based on the material efficiency aspects of personal computers, implementing our strategy would still not be possible. In the future, we may implement a new method (Xing *et al.* 2016) that can achieve a balance between high-speed computation and material efficiency.

Furthermore, only region of interest (ROI) data rather than voxelwise analyses were considered in this study. The characteristics of voxelwise analyses compared to ROI data are different in terms of the between feature correlations and predictor strengths. Previous studies have shown the different results of voxel-based morphometry (VBM) and ROI analyses to detect structural brain alterations between groups (Seyedi, Jafari, Talaei, Naseri, Momennezhad, Moghaddam, and Akbari-Lalimi 2020). Therefore, one could apply the proposed method for VBM analyses in future studies.

### 5.3. Computational time

The high-dimensional problems of these datasets will result in more difficult challenges for the FVS algorithms. The FVS algorithm was applied to analyze the 16S rRNA sequencing microbiome datasets which the number of features was very large (approximately 30,000 features) in our previous study (Dang and Kishino 2022). To reduce the computational burden of the FVS algorithm, some prescreening algorithms (such as the Boruta algorithm (Kursa and Rudnicki 2010) and Laplacian score (He, Cai, and Niyogi 2005)) were proposed to detect all strongly and weakly relevant features to reduce the considerable data dimensionality. With the initial prescreening pipeline, the computational time of the FVS algorithm could be significantly decreased from days to hours (Dang and Kishino 2022). The current pipeline uses a fixed prescreening model. Therefore, additional considerations of this strategy may be necessary if the number of features becomes very large to apply a rigorous search method such as our approach.

## 6. Conclusion

The use of neuroimaging data to train ML models has a significant potential for the identification of brain regions whose structure and activities may contain information predictive of physical phenotypes, mental states and pathological conditions. However, an overwhelmingly large number of ML approaches exist, which may increase the difficulties for those who are unfamiliar with mathematical theories. Moreover, the high-dimensionality of neuroimaging data negatively impacts the power of ML approaches to discover hidden information in the selected neural resources. Furthermore, researchers are often challenged with time-consuming computations to identify neural substrates a variety of neuroscientific discoveries and the development of novel therapeutic interventions. In this study, we proposed a novel procedure that not only automatically selects the best ML model for a specific neuroimaging data, but also identifies a group of brain regions that substantially improve the performance in terms of high-speed computation and high accuracy.

## Acknowledgments

We thank Ms. Haruka Kobayashi for checking source codes in Github.

