## Supplement material for "oFVSD: A Python package of optimized forward variable selection decoder for high-dimensional neuroimaging data"

**Appendix. Description of machine learning algorithms**

In this appendix we provide a theoretical overview for all algorithms evaluated, together with technical details on their implementation.

1. **Ridge regression**

There are response variable $\boldsymbol{y}={\{y_{1}, \ldots, y_{i}\}}_{i=1}^{N}$ and the number of independent features $\boldsymbol{x}= {\{x_{i1}, x_{i2}, \ldots,x_{ip}\}}_{p=1}^{P}$ where N is a number of samples and P is a number of features. The classical way is the ordinary least square (OLS) method which minimizes the squared loss:

$$\sum_{i=1}^{N} {(y_{i}-\sum_{p=1}^{P} \beta_{p}x_{ip})}^{2}$$

(1)

Because the number of features (i.e. number of voxels, cortical vertices or regions) is much larger than the number of samples (p >> n) in high-dimensional datasets, the variance of the estimate *w* by OLS may be large and thus the estimate is not reliable. Ridge regression (Hastie et al., 2009) can reduce the variance by penalizing the norm of the linear transform and minimizes the following cost:

$$J\left( \boldsymbol{\beta} \right)=\sum_{i=1}^{N} {(y_{i}-\sum_{p=1}^{P} \beta_{p}x_{ip})}^{2}+\lambda\sum_{p=1}^{P} \beta_{p}^{2}$$

(2)

Where $\lambda$ is the regularization parameter to controls the trade-off between the bias and variance of the estimate. In practice, one can use cross-validation (Hastie et al., 2009) to find the optimal regularization parameter.

2. **Lasso regression**

Lasso regression replaces squares by absolute values and minimizes the following cost (Hastie et al., 2009; Tibshirani, 1996):

$$J\left( \boldsymbol{\beta} \right)=\sum_{i=1}^{N} {(y_{i}-\sum_{p=1}^{P} \beta_{p}x_{ip})}^{2}+\lambda\sum_{p=1}^{P} \left| \beta_{p} \right|$$

(3)

Ridge regression scales the coefficients by a constant factor, whereas the lasso translates by a constant factor, truncating at 0.

3. **Elastic Net**

The elastic net (EN) method consists of the addition of an L2 penalty to the lasso penalty one to obtain a linear combination of these two norms (Hastie et al., 2009; Zou and Hastie, 2005). The objective is to inherit both the stability of ridge regression, for highly correlated features and the variable selection property of the lasso:

$$J\left( \boldsymbol{\beta} \right)=\sum_{i=1}^{N} {(y_{i}-\sum_{p=1}^{P} \beta_{p}x_{ip})}^{2}+\lambda_{2}\sum_{p=1}^{P} \beta_{p}^{2}+\lambda_{1}\sum_{p=1}^{P} \left| \beta_{p} \right|$$

(4)

4. **Least angle regression (LAR)**

LAR is a computational efficient variant of linear regression with an L1-regularization term (Efron et al., 2004; Hastie et al., 2009). Like the traditional forward selection method, LAR starts with a zero vector as the initial solution (i.e. no active variables), and adds a new predictor variable (i.e. an active variable) at every step. LAR avoids the computational burden of the forward selection method in calculating the coefficients of the active variables.

Specifically, the LAR algorithm includes some main steps:

1. LAR starts with the residual $\boldsymbol{r}=\boldsymbol{y}- \boldsymbol{X\beta}; \beta_{1}= \beta_{2}=\ldots= \beta_{p}=0$.
2. LAR identifies the predictor that correlates most with $\boldsymbol{r}$ (i.e. one that forms the least angle with the residual vector), say $x_{im}$ and add $x_{im}$ to the active set.
3. LAR moves $\beta_{m}$ from 0 towards its least-squares coefficient $\left\langle x_{im},\boldsymbol{r} \right\rangle$ until some other competitor $x_{ik}$ has as much correlation with the current residual.
4. The solution $\beta_{m}$ and $\beta_{k}$ is updated along the direction defined by their joint least squares coefficient of the current residual on $\left( x_{im}, x_{ik} \right)$, until the residuals become equally correlated with another predictor $x_{il}$ which is outside the active set.
5. $x_{il}$ is added to the active set, and the process is repeated until completion or until a desired number of active variables is reached.
5. **The LAR–Lasso combination**

Suppose $\mathbf{A}$ is the active set of variables at some stage in the LAR algorithm, tied in their absolute inner-product with the current residuals $\boldsymbol{r}=\boldsymbol{y}- \boldsymbol{X\beta}$. We can express this as (Hastie et al., 2009)

$$\boldsymbol{x}_{p}^{T} (\boldsymbol{y}- \boldsymbol{X\beta}) = \gamma\boldsymbol{s}_{p}, \forall p \in\boldsymbol{A}$$

Where $\boldsymbol{s}_{p} \in\{-1, 1\}$ indicates the sign of the inner-product, and $\gamma$ is the common value. Now consider lasso criterion in equation 3, $\mathbf{B}$ be the active set of variables in the solution for a given value of $\lambda$ in equation 3. For these variables $J\left( \boldsymbol{\beta} \right)$ is differentiable, and the stationarity conditions give

$$\boldsymbol{x}_{p}^{T} (\boldsymbol{y}- \boldsymbol{X\beta}) = \lambda sign(\beta_{p}), \forall p \in\mathbf{B}$$

6. **Multi-task Lasso regression**

$N_{m}$, P and M denote the number of samples for the m-th task, the number of features for each input matrix, and the number of tasks, respectively. We assume that all the input matrix in $\{X^{m}, m=1, 2,\ldots, M\}$ are having the same dimensionality of features. The multi-task learning with Lasso constraint is given as below (Thung and Wee, 2018; Zhang and Yang, 2022):

$${min \atop\boldsymbol{\beta}}\sum_{m-1}^{M} L\left( \boldsymbol{X}^{m},\boldsymbol{y}^{m},\boldsymbol{\beta}^{m} \right)+ +\lambda\left\| \boldsymbol{\beta} \right\|_{1}$$

(5)

Where $\boldsymbol{\beta=[}\beta^{1}\beta^{2}\boldsymbol{\ldots}\beta^{M}\boldsymbol{]}$ and $\lambda$ is the regularization parameter that controls sparsity in $\boldsymbol{\beta}$

7. **Regularized linear models with stochastic gradient descent (SGD)**

We rewrite the problem in equation 3 of Lasso regression (Shalev-Shwartz and Tewari, 2011):

$\begin{matrix} min \\ \boldsymbol{\beta} \end{matrix}\frac{1}{N} \sum_{i=1}^{N} L\left( \left\langle\boldsymbol{\beta}, \boldsymbol{x}_{i} \right\rangle,y_{i} \right)+ \lambda\left\| \boldsymbol{\beta} \right\|_{1}$

The stochastic coordinate descent algorithm includes main steps:

1. Initialize $\boldsymbol{\beta}$ to be 0
2. At each iteration, we pick a coordinate p uniformly at random from set of features [P] = {1, …, P}
3. The derivative w.r.t the jth feature $g_{j}= \frac{1}{N} \sum_{i=1}^{N} L^{'}\left( \left\langle\boldsymbol{\beta}, \boldsymbol{x}_{i} \right\rangle,y_{i} \right)x_{i,p}$, where L’ is the derivative of the loss function with respect to its first argument.
4. With step size $\boldsymbol{\tau}$, we update $\boldsymbol{\beta}_{p} \leftarrow s_{\lambda/\tau}(\boldsymbol{\beta}_{p}- {g_{j}}/\tau)$
8. **Kernel ridge regression**

Kernel ridge regression combines ridge regression with the kernel trick (Vladimir Vovk, 2013). The data is now replaced with the feature vectors: $x_{i}\to\Phi_{i} = \Phi(x_{i})$ induced by a kernel where $k(x_{i},x_{j}) = {\Phi(x_{i})}^{T}\Phi(x_{j}).$Here $k(.,.)$ is the kernel function which is typically linear $k(x_{i},x_{j}) = {x_{i}}^{T}x_{j}$, polynomial $k(x_{i},x_{j}) ={({x_{i}}^{T}x_{j} + 1)}^{d}$or Gaussian $k(x_{i},x_{j}) = exp(\frac{-\left\| x_{i}-x_{j} \right\|^{2}}{\sigma^{2}})$. Kernel ridge regression minimizes the following cost:

$$J(\boldsymbol{\beta}) = {(\boldsymbol{y} - K\boldsymbol{\beta})}^{T}(\boldsymbol{y} - K\boldsymbol{\beta})+ \lambda\boldsymbol{\beta}^{T}K^{T}\boldsymbol{\beta}$$

Where K is the kernel matrix $K= \left[ \begin{matrix} k(x_{1},x_{1}) & \cdots& k(x_{1},x_{N}) \\ \vdots& \ddots& \vdots\\ k(x_{N},x_{1}) & \cdots& k(x_{N},x_{N}) \end{matrix} \right]$

9. **Decision tree**

A decision tree classifies data items by posing a series of question about the features associated with the items. An internal node contains each question, decision trees are grown by adding question nodes, using labeled training examples to guide the choice of questions. Gini index and entropy are two most common measures that are designed to evaluate the degree of inhomogeneity, or impurity in a set of items. Suppose we want to classify items into K classes. In a node m, representing a region $R_{m}$ with $N_{m}$ observations, the proportion of class k observations in node m is calculated as follow (Hastie et al., 2009):

$$\hat{p_{mk}}= \frac{1}{N_{m}} \sum_{x_{i}\in R_{m}} I(y_{i}=k)$$

Cross-entropy measure is calculated as follow (Hastie et al., 2009):

$$-\sum_{k=1}^{K} \hat{p_{mk}}\log\hat{p_{mk}}$$

where the entropy is lowest when a single $\hat{p_{mk}}$ equals 1 and all others are 0, whereas if all $\hat{p_{mk}}$ are equal, it is the largest.

Gini index is calculated as follow (Hastie et al., 2009):

$$\sum_{k \neq k'} \hat{p_{mk}}\hat{p_{mk'}}=\sum_{k=1}^{K} \hat{p_{mk}} (1- \hat{p_{mk}})$$

To avoid overfitting the training data, we must prune the tree by deleting nodes. There are some approaches such as minimum description length, keep the balance of the complexity of the tree and its fit to the training data by removing internal nodes.

In case of regression, we have p inputs ${\{x_{i1}, x_{i2}, \ldots,x_{ip}\}}_{p=1}^{P}$ and a response $\boldsymbol{y}={\{y_{1}, \ldots, y_{i}\}}_{i=1}^{N}$. we model the response as a constant $c_{m}$ in each region (Hastie et al., 2009):

$$f\left( x \right)= \sum_{m=1}^{M} c_{m} I(x \in R_{m})$$

The greedy algorithm is used to optimate parameters of decision tree regression. There are a number of main steps as follow:

1. Consider a splitting variable j and split point s, and define the pair of half-planes

$R_{1}\left( j,s \right)=\{X|X_{j} \leq s\}$ and $R_{2}\left( j,s \right)=\{X|X_{j} >s\}$

2. For any choice j and s, the inner minimization is solved by

$\hat{c_{1}}= average (y_{i}|x_{i} \in R_{1}\left( j,s \right))$ and $\hat{c_{2}}= average (y_{i}|x_{i} \in R_{2}\left( j,s \right))$

3. The splitting variable j and split point s

$\begin{matrix} min \\ j,s \end{matrix} [\begin{matrix} min \\ c_{1} \end{matrix}\left( y_{i}- \hat{c_{1}} \right)^{2}+ \begin{matrix} min \\ c_{2} \end{matrix}\left( y_{i}- \hat{c_{2}} \right)^{2}]$

10. **Random forest**

In random forest approach, a number of decision trees are grown by a randomized tree-building algorithm. Random forest produces a modified training set of equal size by sampling with replacement the training set. Moreover, this algorithm selects randomly subset of the features when considering the question at each node. There are a number of main steps as follow (Hastie et al., 2009):

1. Consider a number of decision trees B. For each decision tree
   - Draw a bootstrap sample $Z^{*}$ of size N from the training data.
   - Grow a random-forest tree $T_{b}$ to the bootstrapped data, by recursively repeating the following steps for each terminal node of the tree, until the minimum node size is reached.
     i. Select m variables at random from the p variables.
     ii. Pick the best variable/split-point among the m.
     iii. Split the node into two daughter nodes.
2. Output the ensemble of trees ${\{T_{b}\}}_{1}^{B}$

To make a prediction at a new point x:

Regression: $\hat{f}_{rf}^{B}\left( x \right)=\frac{1}{B}\sum_{b=1}^{B} T_{b}(x)$

Classification: $\hat{C_{b}}(x)$ is the class prediction of the bth random-forest tree. $\hat{C}_{rf}^{B}\left( x \right)=majority vote {\{\hat{C_{b}}(x)\}}_{1}^{B}$

11. **Gradient Tree Boosting Algorithm**

Gradient tree boosting algorithm combines multiple decision trees into a stronger algorithm by repeatedly reweighting training examples to focus on the most problematic (Friedman, 2001; Hastie et al., 2009). A constant $\gamma_{j}$ is assigned to each such region and the predictive rule is $x\in R_{j}\Rightarrow f\left( x \right)=\gamma_{j}$. Thus a tree can be formally expressed as

$$T\left( x,\Theta\right)=\sum_{j=1}^{J} \gamma_{j}I(x\in R_{j})$$

with parameters $\Theta={\{R_{j},\gamma_{j}\}}_{1}^{J}$. J is usually treated as a meta-parameter. There are main steps in gradient tree boosting algorithm as follow(Hastie et al., 2009):

1. Initialize $f_{0}\left( x \right)={argmin}_{\gamma}\sum_{i=1}^{N} L(y_{i},\gamma)$
2. For m = 1 to M:
   I. For i = 1, 2, ..., N compute

$$r_{im}=-{[\frac{\partial L(y_{i},f(x_{i}))}{\partial f(x_{i})}]}_{f= f_{m-1}}$$

II. Fit a regression tree to the targets $r_{im}$ giving terminal regions $R_{jm}$, the sizes of each of the constituent trees $J_{m}$, j = 1, 2, ...,$J_{m}$.
III. For j = 1, 2, ...,$J_{m}$ compute

$$\gamma_{jm}=arg\min_{\gamma} \sum_{x_{j}\in R_{jm}} L(y_{i},f_{m-1}\left( x_{i} \right) + \gamma)$$

IV. Update $f_{m}\left( x \right)= f_{m-1}\left( x \right)+ \sum_{j=1}^{J_{m}} \gamma_{jm}I(x\in R_{jm})$

3. Output $\hat{f}\left( x \right)=f_{M}\left( x \right)$
12. **Gaussian Processes**

Multivariate normal distribution is as follow:

$$N\left( x | \mu,\Sigma\right)= \frac{1}{{(2\pi)}^{D/2}\left| \Sigma\right|^{1/2}}exp(-\frac{1}{2}{(x-\mu)}^{T}\Sigma^{-1}(x-\mu))$$

where D is the number of dimensions, x represents the variable, $\mu= E[x]$ is the mean vector, and $\Sigma=cov[x]$ is the covariance matrix. Gaussian kernel function that is defined as

$$cov\left( x_{i},x_{j} \right)=exp(-\frac{{(x_{i} - x_{j})}^{2}}{2})$$

The Gaussian process model is a distribution over functions whose shape is defined by **K**. The standard Gaussian process model is as follow:

$$P\left( \boldsymbol{f} | \boldsymbol{X} \right)=N(\boldsymbol{f}|\boldsymbol{\mu,K})$$

Where the observed data points $\boldsymbol{X=[}x_{1}\boldsymbol{,}\ldots, x_{n}\boldsymbol{]}$**,** $\boldsymbol{f=[}f\boldsymbol{(}x_{1})\boldsymbol{,\ldots,}f\boldsymbol{(}x_{n})\boldsymbol{]}$**,** the mean function $\boldsymbol{\mu=[}m\left( x_{1} \right),\ldots,m\left( x_{n} \right)\boldsymbol{]}$ and positive definite kernel function $K_{ij}=k(x_{i},x_{j})$
